# Breathing new life into the critical oxygen partial pressure (P_crit_): a new definition, interpretation and method of determination

**DOI:** 10.1101/2020.09.01.278440

**Authors:** B. A. Seibel, A. Andres, M. A. Birk, A. L. Burns, C. T. Shaw, A. W. Timpe, C. J. Welsh

**Affiliations:** College of Marine Science, University of South Florida, St. Petersburg, FL 33701; Eugene Bell Center, Marine Biological Laboratory, Woods Hole, MA, 02543

**Keywords:** aerobic scope, hypoxia, metabolic rate, ocean deoxygenation, oxygen and capacity limit of thermal tolerance, oxygen supply, respirometry

## Abstract

The critical oxygen partial pressure (P_crit_) is most commonly defined as the oxygen partial pressure below which an animal’s standard metabolic rate can no longer be maintained. It is widely interpreted as measure of hypoxia tolerance, which influences a species’ aerobic scope and, thus, constrains biogeography. However, both the physiology underlying that interpretation and the methodology used to determine P_crit_ remain topics of active debate. The debate remains unresolved in part because P_crit_, as defined above, is a purely descriptive metric that lacks a clear mechanistic basis. Here we redefine P_crit_ as the PO_2_ at which physiological oxygen supply is maximized and refer to these values, thus determined, as P_crit-*α*_. The oxygen supply capacity (*α*) is a species- and temperature-specific coefficient that describes the slope of the relationship between the maximum achievable metabolic rate and PO_2_. This *α* is easily determined using respirometry and provides a precise and robust estimate of the minimum oxygen pressure required to sustain any metabolic rate. To determine *α*, it is not necessary for an individual animal to maintain a consistent metabolic rate throughout a trial (i.e. regulation) nor for the metabolic rate to show a clear break-point at low PO_2_. We show that P_crit-*α*_ can be determined at any metabolic rate as long as the organisms’ oxygen supply machinery reaches its maximum capacity at some point during the trial. We reanalyze published representative P_crit_ trials for 40 species across five phyla, as well as complete datasets from six additional species, five of which have not previously been published. Values determined using the P_crit-*α*_ method are strongly correlated with P_crit_ values reported in the literature. Advantages of P_crit-*α*_ include: 1) P_crit-*α*_ is directly measured without the need for complex statistics that hinder measurement and interpretation; 2) it makes clear that P_crit_ is a measure of oxygen supply, which does not necessarily reflect hypoxia tolerance; 3) it alleviates many of the methodological constraints inherent in existing methods; 4) it provides a means of predicting the maximum metabolic rate achievable at any PO_2_, 5) P_crit-*α*_ sheds light on the temperature- and size-dependence of oxygen supply and metabolic rate and 6) P_crit-*α*_ can be determined with greater precision than traditional P_crit_.

## Introduction

The relationship between metabolic rate and environmental oxygen has long been of interest due to its implications for human health, paleobiology, fisheries management, biogeography, species diversity and evolution (Tang, 1933; Hall, 1966; Fry and Hart, 1948; Seibel and Deutsch, 2020; Penn et al., 2018; Childress and Seibel, 1998; Deutsch et al., 2015; 2020; Ern et al., 2016; Rogers et al., 2016; Ultsch and Regan, 2019). This relationship has taken on special significance in light of anthropogenic ocean warming and deoxygenation (Seibel and Wishner, 2019; Breitburg et al., 2018). The critical oxygen partial pressure (critical PO_2_, P_crit_) is a metric commonly interpreted as a measure of hypoxia tolerance, with a lower P_crit_ indicating greater tolerance for low oxygen (e.g. Rogers et al., 2016; Seibel, 2011). The ratio of environmental PO_2_ to P_crit_, normalized for temperature, provides a direct measure of aerobic scope that is believed to constrain the viability of current and future habitat (Deutsch et al., 2015; Deutsch et al., 2020). However, hypoxia and tolerance are subjective, time-dependent terms while P_crit_ is inconsistently defined and determined, resulting in debate about its meaning and significance (Wood, 2018; Regan et al., 2019).

An organism’s P_crit_ is typically defined based on the physiological response to declining environmental PO_2_, at a clear decline (breakpoint) in the measured metric at the P_crit_ (Pörtner and Grieshaber, 1993; Richards, 2011; Hall, 1966; Ultsch and Regan, 2019; Childress and Seibel, 1998). Metrics employed to determine critical oxygen levels include cardiac output, anaerobic metabolite concentrations, gene expression, and behavioral impairment (e.g. loss of equilibrium) (Mandic et al., 2009; Ultsch and Regan, 2019; Farrell and Richards, 2009 for reviews). However, the most commonly measured response variable is the oxygen consumption (metabolic) rate, with the corresponding definition of P_crit_ as the minimum PO_2_ at which a given metabolic rate can be sustained. Accordingly, the rate of oxygen consumption becomes dependent on PO_2_ below P_crit_. Because of the rate dependence of P_crit_, as typically defined, recent protocols recommend measuring P_crit_ at the standard metabolic rate (SMR; Reemeyer and Rees, 2019). At P_crit_ for SMR, by definition, metabolism is oxygen limited and aerobic scope (the difference between resting and maximum rates) is abolished. This is what Fry and Hart (1948) referred to as the “oxygen level of no excess activity” or the “incipient lethal oxygen level”. In order to determine P_crit_ using this definition, the resting metabolic rate must be established and, preferably, maintained at high PO_2_ (i.e. it must be “regulated”). A break-point must be readily apparent between oxygen dependent and oxygen independent portions of the respiration curve. Alternatively, complex statistics may be used to identify an inflection point in the data (Muggeo, 2003).

At PO_2_ levels above P_crit_, a constant oxygen consumption rate (MO_2_ or VO_2_ in moles or volume of oxygen, respectively) is typically considered to be “regulated” and independent of PO_2_. Current P_crit_ methodology places a great deal of importance on this constancy and the degree of oxygen independence of the regulated rate at high PO_2_ values. In fact, several methods have been developed to quantify oxyregulation, loosely defined as the overall respiratory response of organisms over the complete range of measured oxygen pressures (e.g. *Regulation Index*, Mueller and Seymour, 2011; Tang, 1933; non-linear regression, Marshall et al., 2013; and best-fit approaches, Cobbs and Alexander, 2018; Muggeo, 2003). Proponents of these methods argue that a great deal of information is lost by distilling a respirometry trial down to a single critical PO_2_ (Marshall et al., 2013; Wood, 2019), although it is unclear whether analyzing the relationship between MO_2_ and PO_2_ at PO_2_ greater than P_crit_ provides any additional information. More importantly, any information provided by P_crit_ is lost in mathematical descriptions of the entire trial. The term “regulation” has a specific implication that has injected a great deal of confusion into our collective understanding of P_crit_, and, thus, how it should be measured. The concept of regulation, as applied to other physiological processes, implies that an organism strives for homeostasis by maintaining the constancy of internal conditions (e.g. acid-base or osmotic balance; Trowell, 1943). However, a metabolic rate is merely a measure of the energy demand at a point in time and, although there are upper and lower bounds on the rate of metabolism and the rate is under active control, it is not regulated to achieve homeostasis. There is no specific level of metabolic cost that must be maintained for proper function.

Unless the relationship between MO_2_ and PO_2_ is causal, there is no particular reason to describe it at all. At PO_2_ above P_crit_, MO_2_ may correlate with a number of covariables, including time in captivity, time since feeding, accumulation of metabolic waste, diel cycles, light, stress and activity. Some of these variables may be controlled for during experiments, but even so, as Ultsch and Regan (2019) point out, it is impossible to know whether MO_2_ above P_crit_ is supporting the same maintenance processes that comprise SMR at P_crit_: *“When PO*_*2*_ *is reduced, there may be (and probably is) a reallocation of resources, including O*_*2*_, *towards a different suite of processes (e.g. ventilation) that may nevertheless sum to a value of ṀO*_*2*_ *similar to normoxic SMR”* (Ultsch and Regan, 2019). Thus, apparent oxyregulation is not necessarily oxygen independence and a lack of apparent regulation (i.e. oxyconformation) is not necessarily oxygen dependence. Rather, what we typically refer to as oxyregulation are physiological adjustments that provide additional oxygen as environmental PO_2_ declines, regardless of the constancy or level of metabolic rate.

The plethora of incompatible and inconsistent definitions of, and methods for, describing the response of metabolism to declining PO_2_ underscores an incomplete physiological understanding. All existing definitions focus on consequences, rather than causes, of physiological failure at low oxygen. The lack of a specific mechanistic basis limits the generality and utility of P_crit_ (Wood, 2018). Although oxygen supply limitation below a critical PO_2_ is implied, oxygen supply itself is rarely determined. Here we redefine P_crit_ as the PO_2_ at which physiological oxygen supply mechanisms are operating at maximum capacity (Deutsch et al., 2015; 2020; Seibel and Deutsch, 2020; Kielland et al., 2019). Values thus determined, designated P_crit-*α*_, are not specific to any particular metabolic rate (MR) and can be mathematically defined as P_crit-*α*_ = *MR/α*, where *α* is the maximum physiological capacity for oxygen supply (see detailed derivation below). This straightforward definition provides a strong theoretical justification for the physiological value of P_crit_ and eliminates most, if not all, of the recently described pitfalls associated with its measurement (Wood, 2018; Reemeyer and Rees, 2019; Ultsch and Regan, 2019; Regan et al., 2019). This new definition also provides some clarity to the concept and calls into question widely held beliefs about the ecological and evolutionary significance of P_crit_. As stated by Fry and Hart (1948), “*the worth of such data to the ecologist must ultimately depend on proof that they have real significance as values limiting the activity of the organism in nature*.” Many decades later such proof is still not readily available and most evidence is merely correlative. Strong comparative approaches have been employed to demonstrate that P_crit_ sometimes reflects physiological adaptations to low oxygen (Regan et al., 2019; Wishner et al., 2018; Childress and Seibel, 1998; Mandic et al., 2009), but the ecological relevance of a particular P_crit_ value is often assumed. More troubling is that P_crit_ is often used to infer hypoxia tolerance in the absence of any measure of ambient PO_2_ in the environment.

Seibel and Deutsch (2020) argued that P_crit_ is not a measure of hypoxia tolerance, but rather a measure of the oxygen supply capacity, which evolves to support maximum metabolic rate at the prevailing (highest persistently encountered) environmental PO_2_ (Fig. 1). Because P_crit-*α*_ provides a measure of oxygen supply capacity, it can be used to calculate MMR and aerobic scope (Seibel and Deutsch, 2020; Deutsch et al., 2020; Howard et al., 2020). In other words, the ratio of MR:PO_2_ should reach a maximum at P_crit-*α*_ regardless of the metabolic rate. This ratio provides a simple, direct, and quantitative method of determining *α* that does not require an obvious break-point in, nor a particular level of, metabolic rate.

**Figure 1.**
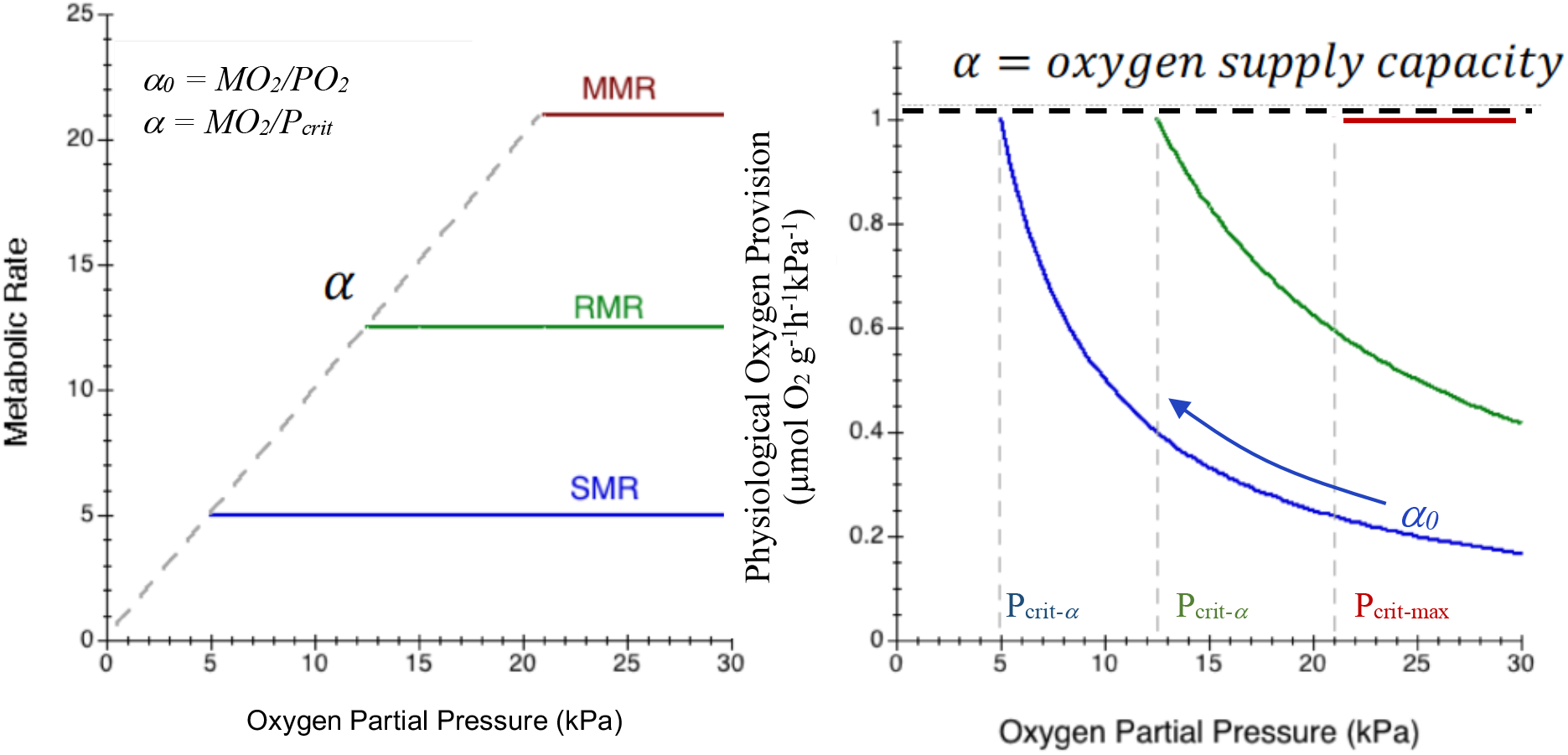
Schematic illustrating a new method for P_crit_ determination, termed P_crit-*α*_. A) The relationship between standard (SMR, blue), routine (RMR, green) and maximum (MMR, red) metabolic rate and oxygen partial pressure (PO_2_). The oxygen supply capacity (*α*, dashed line, derived in panel B) is the slope of the relationship between P_crit-*α*_ and PO_2_. B) For each metabolic level (e.g. SMR), the physiological oxygen provision (*α*_*0*_) increases toward the maximum capacity, *α* (dashed line). While *α*_*0*_ must be greater to meet a higher metabolic rate at a given PO_2_, *α* is the same for all three rates. All three metabolic levels reach a critical PO_2_ where oxygen provision is at maximum capacity (where *α*_*0*_ *= α*). As a result, *α* is the slope of the MO_2_ vs PO_2_ curve, allowing P_crit_ to be calculated at any metabolic rate and the maximum achievable metabolic rate to be calculated for any PO_2_.

### Methods: Oxygen supply capacity and P_crit-*α*_ determination

For an aerobic organism to obtain sufficient energy for survival, its oxygen supply must meet its demand. Oxygen demand is the metabolic rate (MR), most commonly measured at rest (standard, SMR) or at maximum exertion (MMR, the maximum achievable rate for a given PO_2_). The total amount of oxygen available for cellular respiration is a function of the ambient environmental PO_2_ and the physiological oxygen provision, *α*_*0*_ (Seibel and Deutsch, 2020; Deutsch et al., 2015). Physiological oxygen provision comprises each step in the oxygen cascade (Weibel et al., 1991) from ventilation and blood-oxygen binding to cardiac output and circulation. At a constant metabolic rate, physiological oxygen provision must increase as environmental PO_2_ declines (Fig. 1). We define and calculate *α*_*0*_ as the rate of oxygen consumption per unit of available environmental oxygen pressure (*α*_*0*_ = MR/PO_2_; µmol O_2_ g^-1^h^-1^kPa^-1^). The P_crit-*α*_ is reached when physiological oxygen provision is operating at its maximum capacity (*α*; Fig. 1; *Eq. 1*). At capacity, these functional traits cannot be further upregulated and the metabolic rate is entirely dependent on, and directly proportional to, environmental PO_2_. Thus, the physiological oxygen supply capacity (*α* = MR/ P_crit-*α;*_ Fig. 1) is set by attributes of the oxygen cascade that are specific to each species and to acclimation conditions. As a result, *α* describes the slope of the relationship between MMR and PO_2_ and can be used to calculate P_crit-*α*_ at any metabolic rate (P_crit-*α*_ = MR/*α*). By rearranging this equation (MMR = *α\**PO_2_), *α* can also be used to calculate MMR at any PO_2_ up to P_crit-max_ (the PO_2_ above which no additional increase in MO_2_ is possible, even in hyperoxia; Seibel and Deutsch, 2020). Thus, P_crit-*α*_ is defined by maximized oxygen supply, rather than the oxygen dependence of a particular metabolic rate.

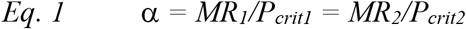

To assist in the adoption of this new method, calculation of *α* and P_crit-*α*_ have been incorporated into the R package “respirometry” with an optional parameter available to define the metabolic rate of interest for P_crit_ estimation (Birk 2020). The “calc_pcrit” and “plot_pcrit” functions now return side-by-side comparisons of P_crit_ calculated using a broken-stick regression (Yeager and Ultsch 1989), nonlinear regression (Marshall et al 2013), LLO method (Reemeyer and Rees, 2019); sub-prediction interval method (Birk et al. 2019), and the P_crit-*α*_ method presented here. This software is freely available on CRAN at https://cran.r-project.org/package=respirometry.

The oxygen supply capacity (*α*) has been recently calculated from P_crit_, determined using various historical methods, and from the metabolic rate at which it was determined (Deutsch et al., 2015; Seibel and Deutsch, 2020; Kielland et al., 2019). Here we show that *α* can be directly measured as the highest *α*_*0*_ achieved during a respirometry trial and used to determine P_crit_ (P_crit-*α*_). P_crit-*α*_ is then calculated from *Eq. 1* for every metabolic rate bin measured during a trial, providing a straight line that identifies P_crit-*α*_ for *any* metabolic rate (Fig. 2). We used several approaches to test the applicability, generality and precision of this novel P_crit-*α*_ method. We first extracted and reevaluated available literature data for 40 species in five phyla (Table 1). Most studies that report P_crit_ provide only summarized data in the text but some studies included a “representative” trace showing the relationship between the oxygen consumption rate and PO_2_. Data were extracted from these respirometry curves using WebPlotDigitizer 4.2 (Rohatgi, 2019; https://automeris.io/WebPlotDigitizer/index.html). The oxygen supply capacity was determined and used to calculate P_crit-*α*_ across the range of measured metabolic rates. We suspect that these representative curves are likely the best of many trials, rather than being truly representative. Thus, they may not provide the most rigorous test of the new method. Accordingly, we also tested our method on existing datasets for zooplankton (Wishner et al., 2018), shrimp (Burns et al., unpubl.), squids (Birk and Seibel, unpubl.) and fishes (Slesinger et al., 2019; Andres et al., submitted). We tested inter-individual and temperature-related variances in *α* as well as the effect of duration bin size when calculating MO_2_.

**Table 1.**
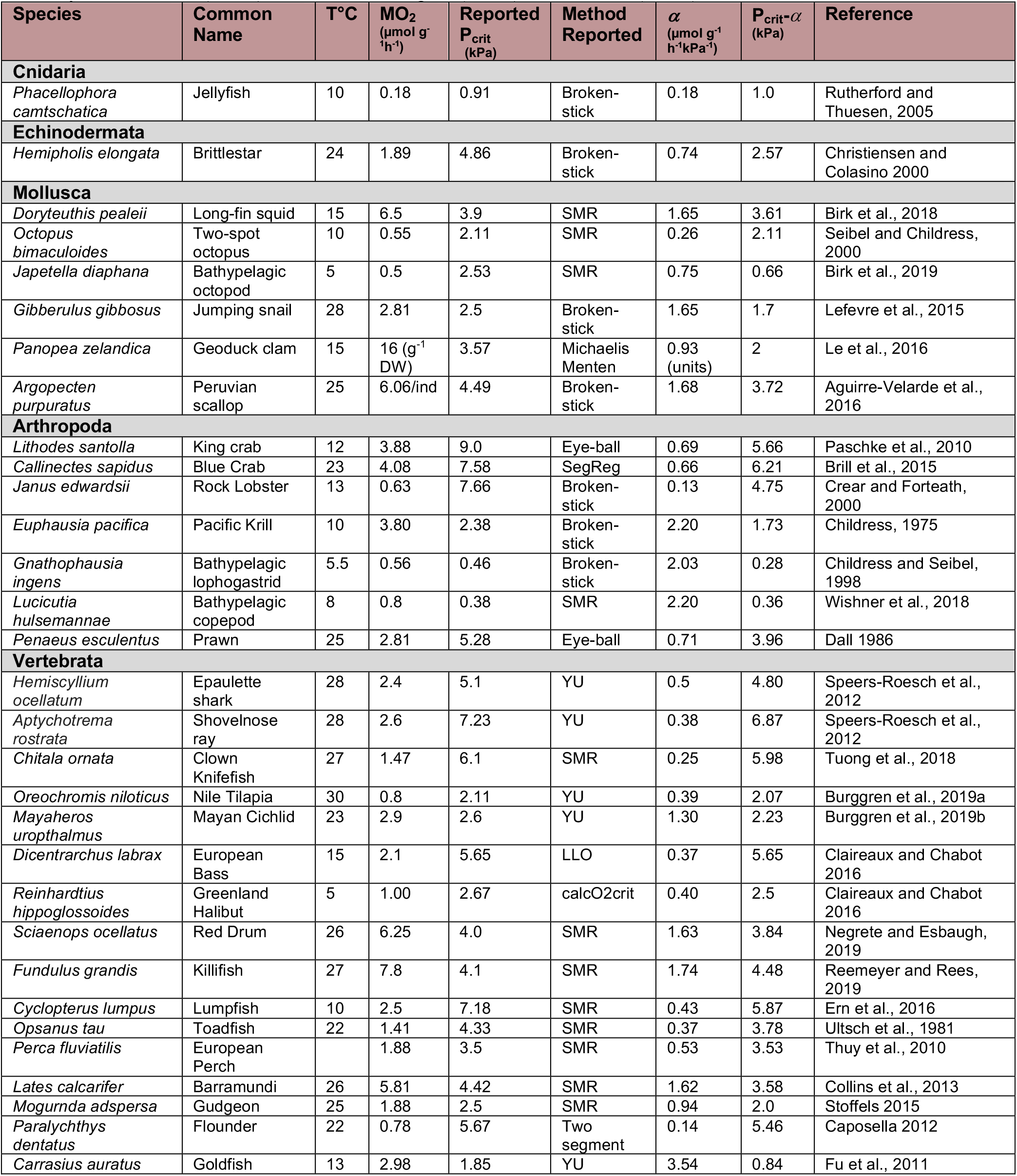
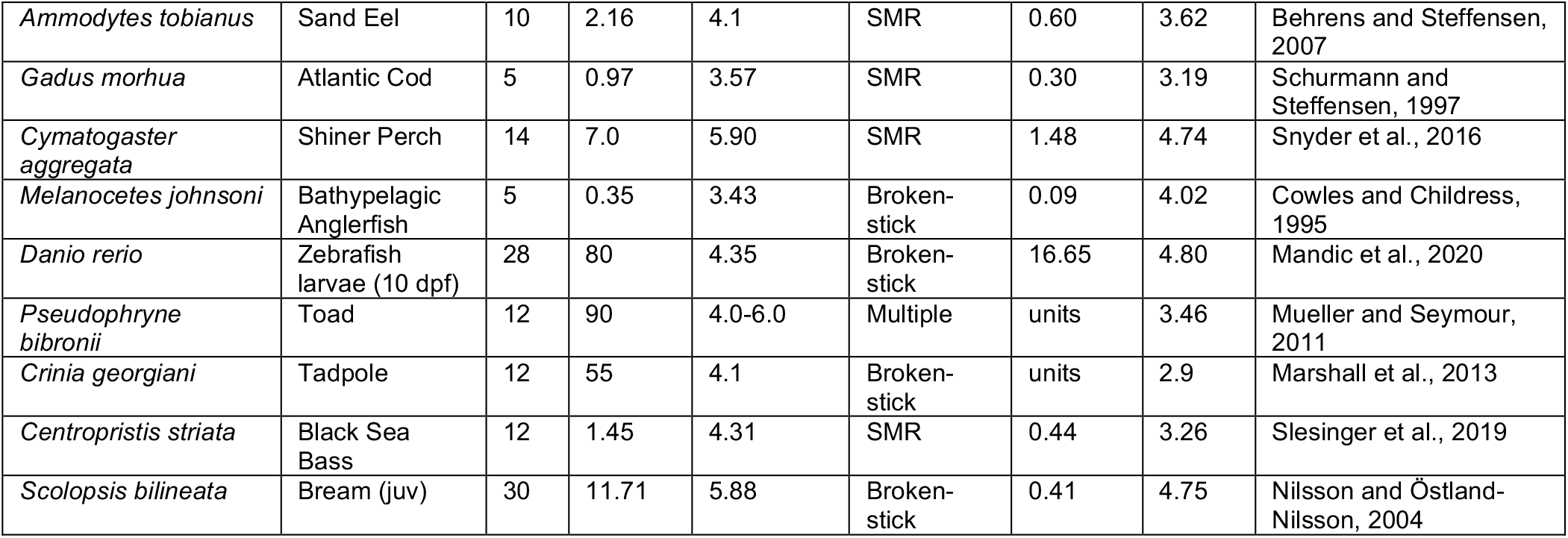
P_crit-*α*_ compared to published P_crit_ measurements for diverse aquatic animals. The methods used for published Pcrit estimates include those described by YU (Yeager and Ultsch, 1989), LLO (Reemeyer and Rees, 2019), and SMR (Chabot and Claireaux, 2008). See Reemeyer and Rees (2019), Ultsch and Regan (2019) and Wood (2018) for recent reviews.

**Figure 2.**
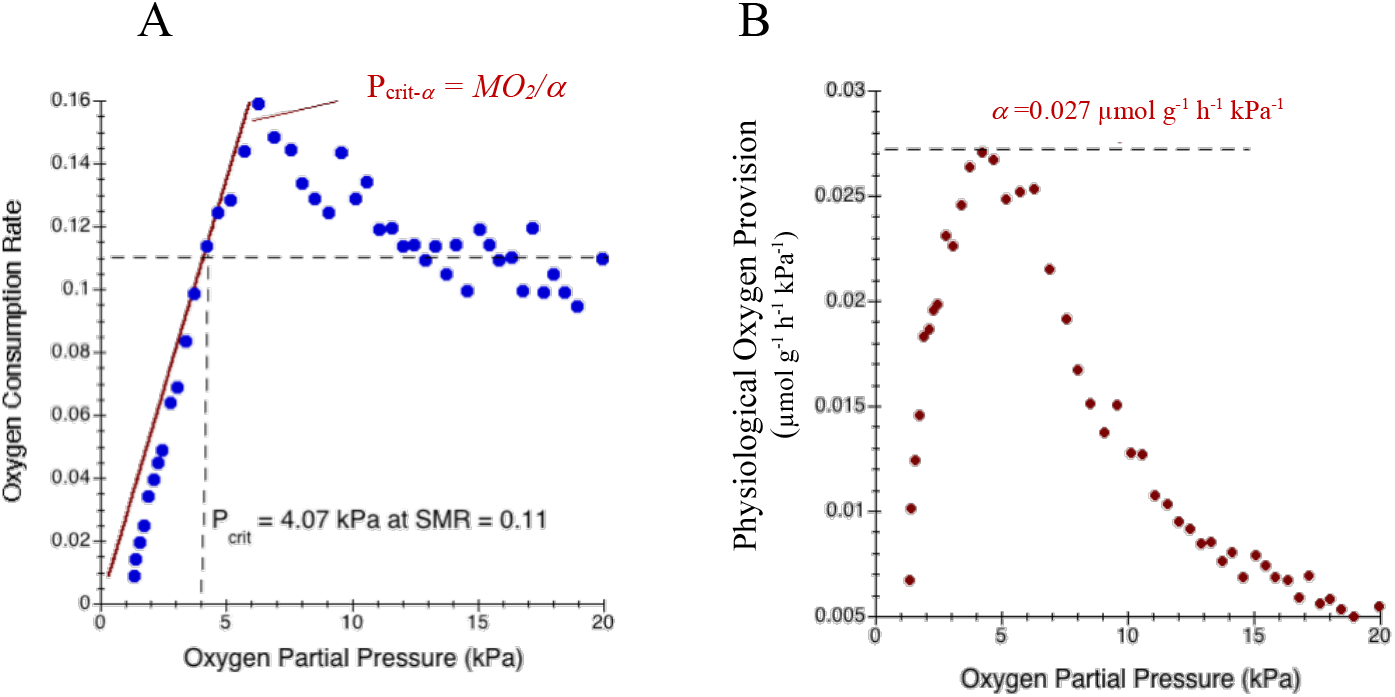
A new method for P_crit_ determination using a representative respirometry trial for *Fundulus grandis* (Reemeyer and Rees, 2019). A) Oxygen consumption rate (MO_2_, blue dots) as a function of PO_2_. P_crit-*α*_ (red line; P_crit-*α*_ = MO_2_*/α*) is the MO_2_ value divided by the oxygen supply capacity, *α* (derived in panel B). Thus, P_crit-*α*_ can be determined for any MR. B) Each MO_2_ bin (panel A) divided by its corresponding PO_2_ is the physiological oxygen provision (*α*_*0*_; µmol O_2_ g^-1^h^-1^kPa^-1^). As PO_2_ declines, oxygen provision increases toward the oxygen supply capacity (*α* = 0.027, dashed line). Using several common methods for P_crit_ determination, including the traditional broken-stick method, non-linear statistics, and a segmented approach with independent SMR determination, P_crit_ for this trial varied from 2.9 to 9.9 (Reemeyer and Rees, 2019).

## Results

Table 1 presents P_crit-*α*_ and published P_crit_ for representative traces determined at the metabolic rate provided in the literature (n = 40). For comparative purposes, the rates and PO_2_ values are converted to common units in Table 1 and in the text. In the species analyzed here, *α* ranged from 0.09 µmol O_2_ g^-1^h^-1^kPa^-1^ for the deep-sea anglerfish, *Melanocetus johnsoni*, at 5°C (Cowles and Childress, 1995), to 16.65 µmol O_2_ g^-1^h^-1^kPa^-1^ at 28°C for larval zebrafish, *Danio rerio* (Mandic et al. 2020; Table 1). These values are within the previously reported range and are correlated with metabolic rate (Deutsch et al., 2020; Seibel and Deutsch, 2020; Kielland et al., 2019). P_crit-*α*_ was strongly correlated with published P_crit_ values that were determined using a variety of methods (P_crit-*α*_ = 0.14 + 0.80P_crit_; R^2^= 0.83; n = 40; Fig. 3). For a subset of the trials extracted from the literature, SMR and P_crit_ were determined as in Chabot and Claireaux (2008) and Reemeyer and Rees (2019), respectively. For these cases, P_crit-*α*_ and the reported P_crit_ were more tightly correlated because the metabolic rates used to determine P_crit_ were more tightly constrained (y = 0.29 + 1X; R^2^ = 0.99; n =6; Fig. 3A).

**Figure 3.**
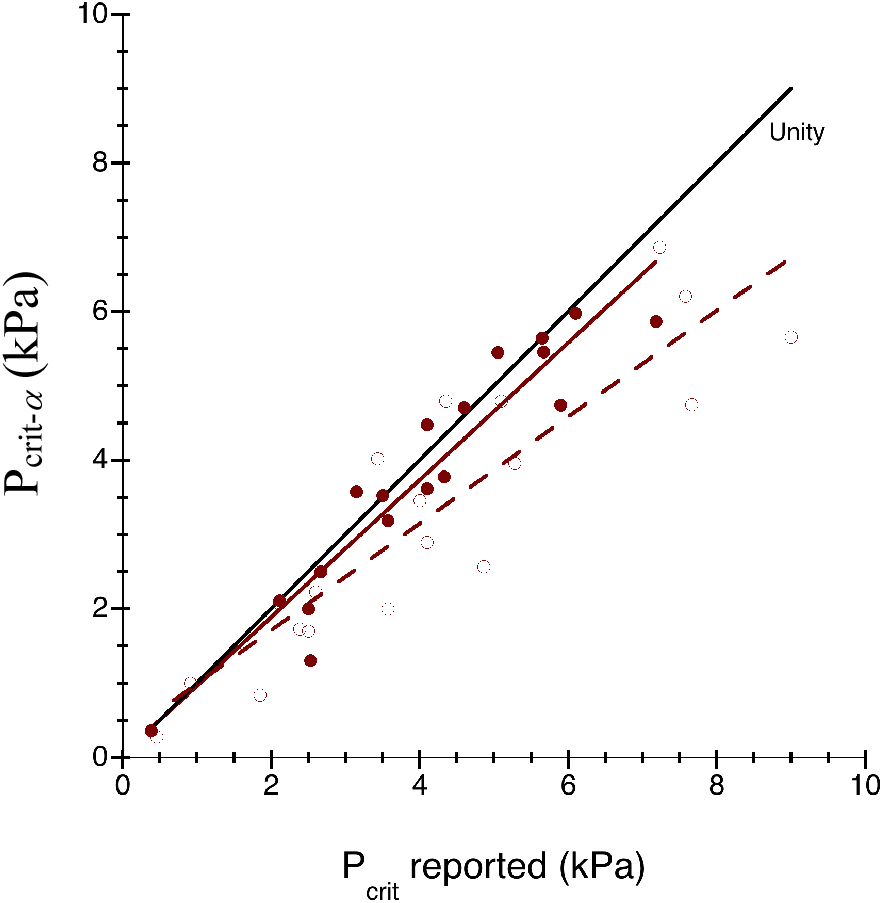
Critical oxygen partial pressure (P_crit-*α*_, kPa), determined using the present method, as a function of that determined using other published methods (P_crit_ reported) at an equivalent metabolic rate. P_crit_ measured using the LLO method (Reemeyer and Rees, 2018; closed symbols, solid line) is most strongly correlated (P_crit-*α*_ = 0.046 + 0.92P_crit-reported_; R^2^ = 0.90). Open symbols (dashed line) were measured with a variety of other methods (Table 1; P_crit-*α*_ = 0.27 + 0.72 P_crit-reported_; R^2^ = 0.81). The black line is a unity line indicating a 1:1 relationship.

The physiological oxygen provision (*α*_*0*_ =MO_2_/PO_2_) was calculated for each MO_2_ value from all available trials. In our dataset, *α*_*0*_ responded to declining PO_2_ in one of three distinct ways; a peak (with subsequent decline), a plateau, or a continuous increase. In most cases, *α*_0_ increased to a peak or plateau as PO_2_ decreased (Fig. 4). A peak in *α*_0_, with a subsequent decline at lower PO_2_, indicates that the individual is failing physiologically and MO_2_ is declining faster than PO_2_ below P_crit_. A plateau in *α*_0_ indicates maintenance of oxygen supply capacity as PO_2_ declines linearly below P_crit_ toward anoxia. In either case, the highest resulting *α*_*0*_ value is the oxygen supply capacity, *α*, which was used to calculate P_crit-*α*_ across the range of MO_2_ values in each trial (P_crit-*α*_ = MR/*α*). A continuous increase in *α*_*0*_ (Fig. 4C), with the highest value occurring at the lowest PO_2_, provides no clear peak or plateau and renders *α* more dubious. This represents either a decline in MO_2_ before P_crit_ is reached or possible error in PO_2_ measurement (see below). A small number of published representative trials (2 of 40) exhibited apparent “oxyconformation”, where MO_2_ declined continuously with PO_2_ with no obviouis breakpoint (Fig. 5). For such trials, *α*_*0*_ was either independent of PO_2_, suggesting that MO_2_ is maximized throughout the experiment (Fig. 5A), or it increased continuously as PO_2_ declined, never reaching a peak or plateau (Fig. 5B).

**Figure 4.**
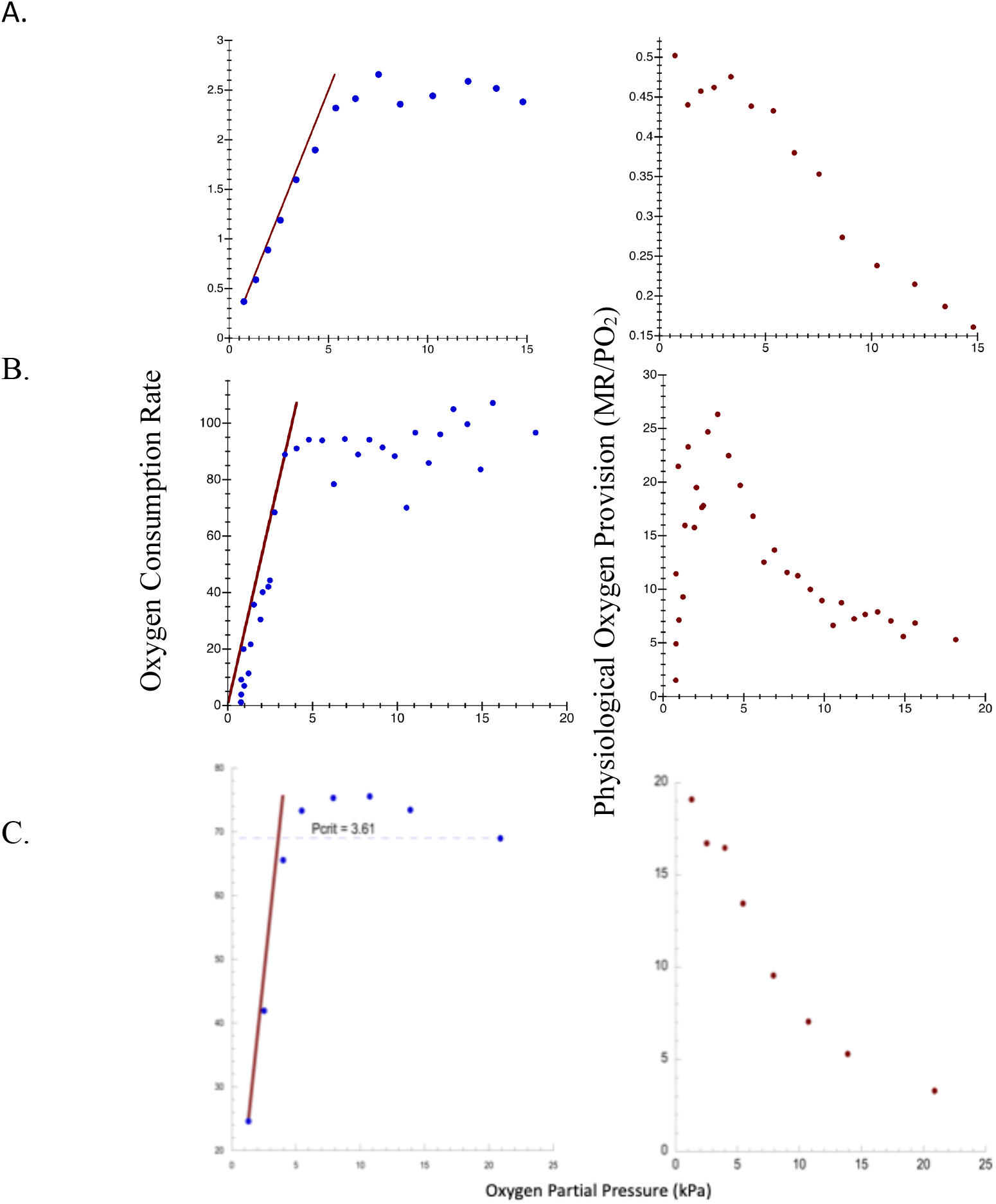
Three common response types of physiological oxygen provision (*α*_*0*_) vs oxygen partial pressure. In each pair (A, B and C) the left-hand panel shows the oxygen consumption rate (blue dots) as a function of PO_2_, with P_crit-*α*_ for each rate indicated by the red line (P_crit-*α*_ =MO_2_/*α*). The oxygen supply capacity (*α*) is derived in the right panel. The physiological oxygen provision (*α*_*0*_ = MO_2_/PO_2_) increases toward capacity (*α*) as PO_2_ declines. A) Epaulette shark, *Hemiscyllium ocellatum* (Speers-Roesch et al., 2012). The oxygen provision plateaus at P_crit_, meaning that oxygen supply is maintained at full capacity until oxygen is exhausted. B) *Pseudophryne bibronii*, post-hatch (Mueller and Seymour, 2011). Oxygen provision peaks at P_crit_ and declines at lower PO_2_, meaning that the animal is failing at PO_2_ less than P_crit_. C) Sand eel, *Ammodytes tobianus* (Behrens and Steffensen, 2007). Oxygen provision peaks at the lowest PO_2_, which suggests that MO_2_ begins to decline just before P_crit_ is reached. Note that all MO_2_ and *α* values in this figure are expressed in the units as originally published.

**Figure 5.**
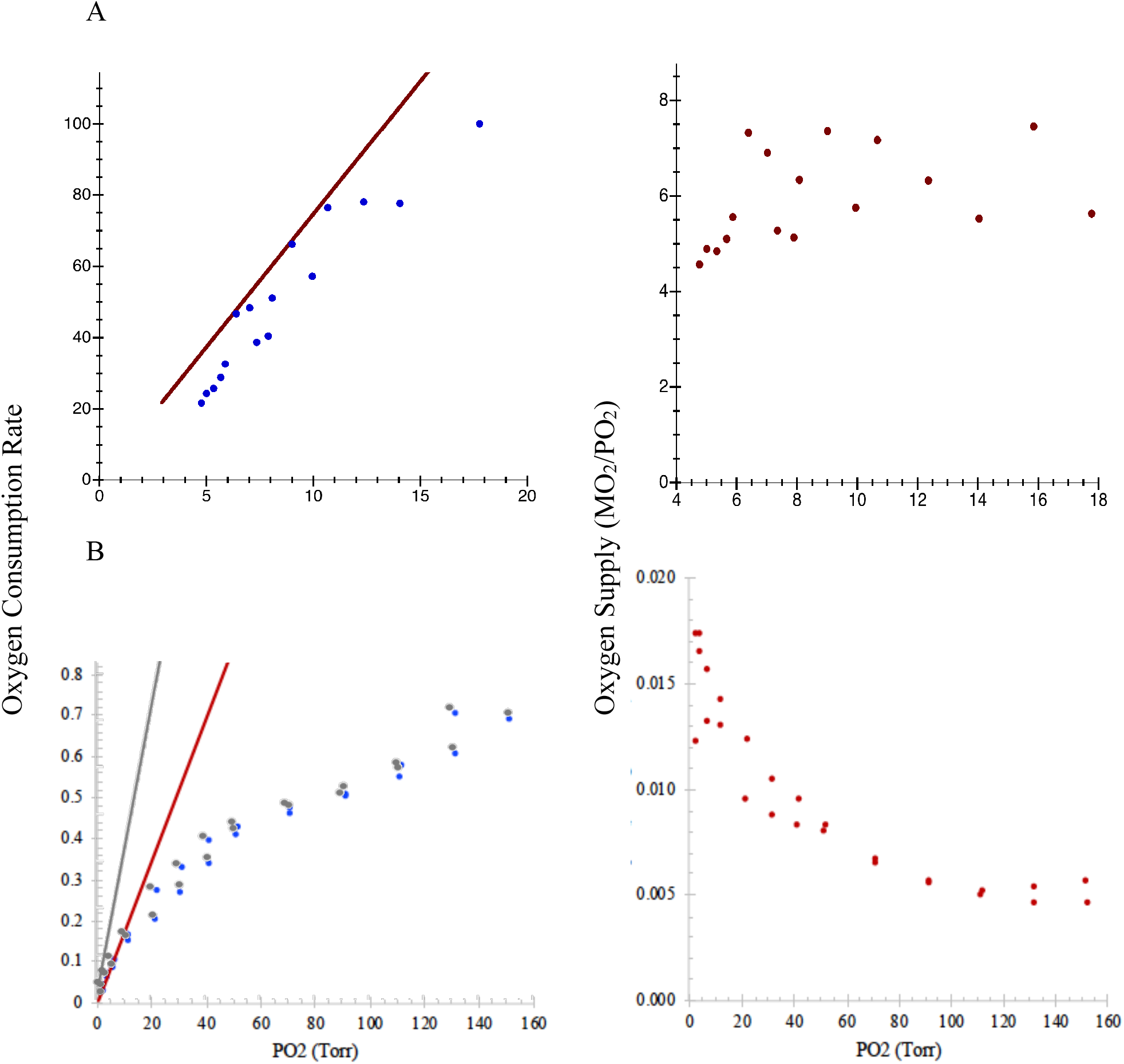
Apparent oxyconformation in A) the clown knifefish, *Chitala ornata*, and B) in the worm, *Sipunculus nudus*. In the knifefish (*C. ornata*, 33°C), oxygen supply (right, red dots) is always near capacity and does not change appreciably as PO_2_ declines, indicating that this individual was operating near the maximum achievable metabolic rate throughout the trial. B) The worm, *S. nudus*, oxygen supply increases as PO_2_ and MO_2_ decline, suggesting some active regulation despite apparent oxyconformation. The *S. nudus* data were corrected for PO_2_ error (Timpe et al., in prep; gray points in B), which led to a reduction in *α* from 0.052 to 0.017 (gray and red lines). As a result, the ratio of maximum to minimum *α*_0_ (a measure of the factorial aerobic scope) was reduced from 11.3 to 3.8, which is within the range of most species. Units are as in the original publications.

To assess intraspecific variability and to ensure that our analysis was not biased by the use of idealized representative curves from the literature, we also analyzed several existing respirometry datasets-*Euphausia mucronata*, a midwater krill from the Eastern Tropical Pacific oxygen minimum zone (OMZ; Welsh and Seibel, unpublished), *Farfantepenaeus duorarum*, an estuarine pink shrimp (Burns et al., unpublished), the Atlantic Spiny Dogfish, *Squalus acanthias* (Andres et al., submitted), two oceanic squids, *Illex illecebrosus* and *Sthenoteuthis oualaniensis* (Birk, unpublished), and the Black Seabass, *Centripristis striata* (Slesinger et al., 2019), a coastal fish species. The mean values of *α* and P_crit-*α*_ and P_crit_ (LLO or breakpoint method) for these species are shown in Table 2. The values of *α* were highest in the squids, due to their high metabolic rates, and in the crustaceans, due to high rates at small sizes and, likely, adaptation to their hypoxic OMZ and estuarine habitats (Table 2). With the exception of crustaceans from hypoxic environments, P_crit-*α*_ was very consistent at a given temperature (ranging from 4.04 to 5.18 at ∼20-23°C, n = 4 species). The *α*_*0*_ response types (see Fig. 4) varied among individuals of each species, but typically displayed a clear peak or plateau, facilitating identification of a clear *α*. The length of respirometry measurement periods, or “bin sizes”, had an effect on *α*. Small bin sizes (< 5min) gave undue influence to outliers among the MO_2_ measurements leading to unrealistically high *α* and low P_crit-*α*_. With increasing bin size, *α* declined and was consistent between bin sizes of 10, 15 or 20 min (Fig. 6). For krill, smaller bin sizes similarly resulted in a slight increase in *α* but did not have a significant effect on P_crit-*α*_. The pattern for krill was similar to the literature values mentioned above, with Pcrit slightly, but consistently, higher than P_crit-*α*_. For Atlantic Spiny Dogfish and Black Seabass, P_crit-*α*_ matched P_crit_ determined using the LLO method (Reemeyer and Rees, 2019) across a range of temperatures and *α* increased with temperature in proportion to MMR (Fig. 6; Slesinger et al., 2019; Andres et al., in prep).

**Table 2.**
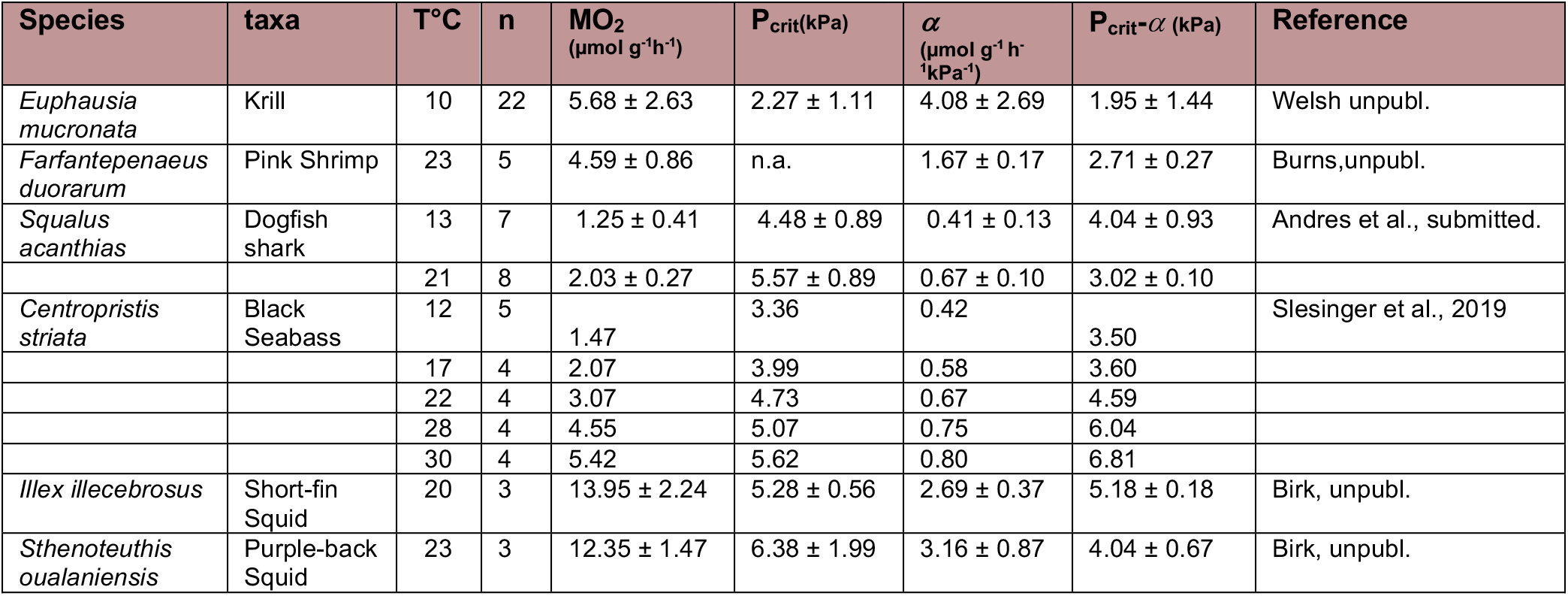
P_crit-*α*_ compared to P_crit_ derived using LLO (Reemeyer and Rees, 2019) or broken-stick analyses (Yeager and Ultsch, 1989) for complete datasets to assess intraspecific variability.

**Figure 6.**
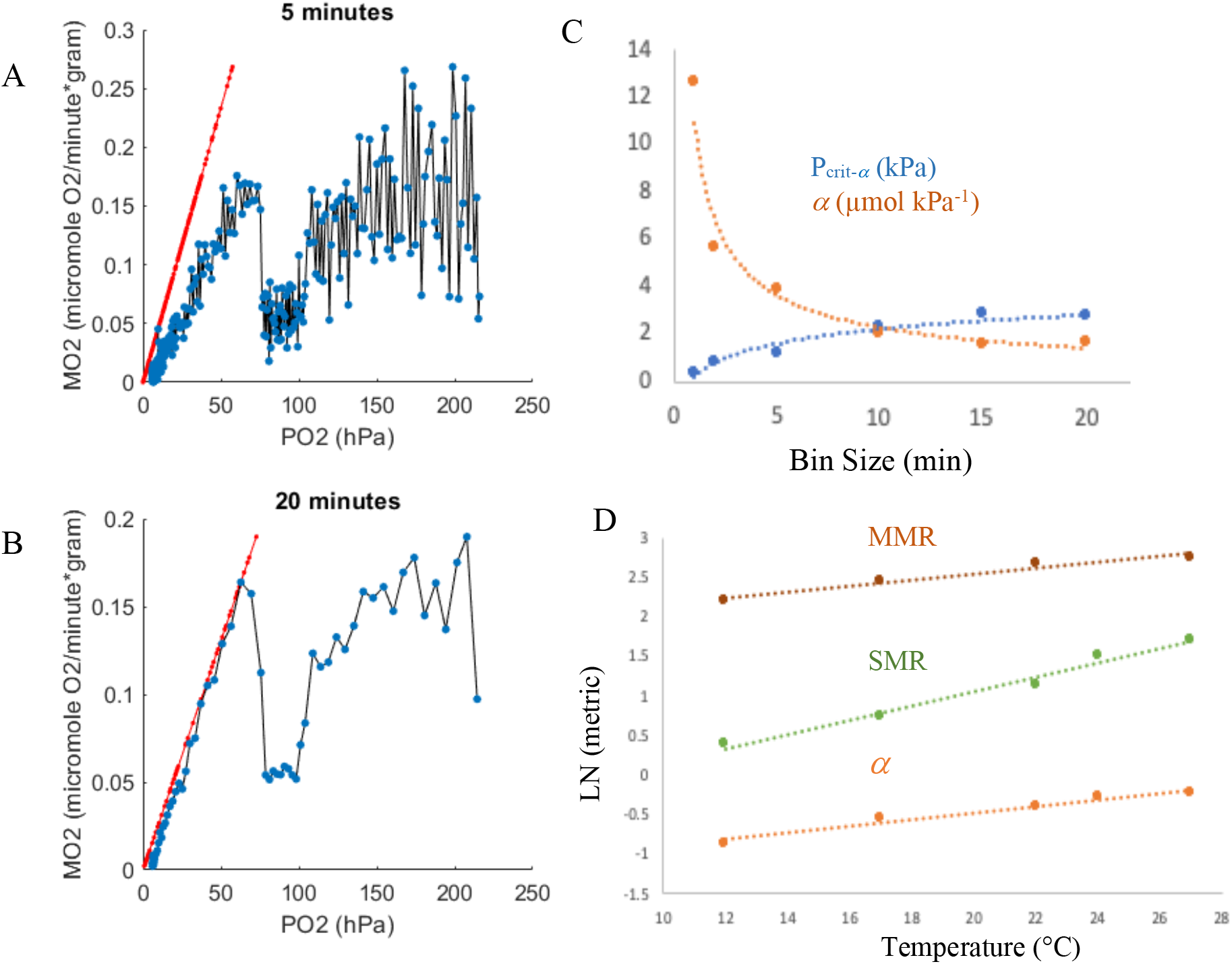
Temperature, methodological, and inter-individual variation in P_crit-*α*_ for existing datasets. A-C) Shrimp, *Farfantepenaeus duorarum* (Burns, unpubl.): Oxygen consumption rate as a function of PO2 for an individual animal was calculated over time bins varying from 1 to 20 minutes. The 5- (A) and 20-min (B) time bins are shown. Oxygen supply capacity (*α*, slope of the red P_crit-*α*_ line) was strongly influenced by increasing MO_2_ outliers at small bin sizes. C) High *α* (orange) at small bin size lead to unrealistically low P_crit-*α*_ (blue). No significant differences were found among values obtained using 10 to 20 min bin sizes. Trials were ∼12 h duration and oxygen was recorded every 30 s. D) Black Seabass (*Centropristis striata*, Slesinger et al., 2019): Oxygen supply capacity (*α*) increases with temperature in proportion to maximum (MMR), rather than standard (SMR) metabolic rate. B)

P_crit-*α*_ was typically slightly lower than the reported P_crit_ for an equivalent metabolic rate, in part because P_crit-*α*_ is based on the highest calculated oxygen supply, *α*_*0*_, which places P_crit-*α*_ (the oxygen supply limit; red lines in Fig 2E and Figs. 4-7) on or to the left of all MO_2_ values. This precludes the possibility that PO_2_ is underestimated due to measurement error, which, of course, is not a valid assumption. However, previous P_crit_ methods assume that PO_2_ error is spread evenly around the PO_2_ vs MO_2_ regression line (the oxygen-dependent portion of the MO_2_ trial) with a Gaussian distribution. This assumption is problematic if error is minimal (i.e. if scatter in the data is real variation in the organism’s oxygen consumption rate) because it suggests that oxygen is sometimes consumed faster than it is supplied. Regardless, P_crit-*α*_ and P_crit_ derived from other methods were not significantly different (Fig. 3).

**Figure 7.**
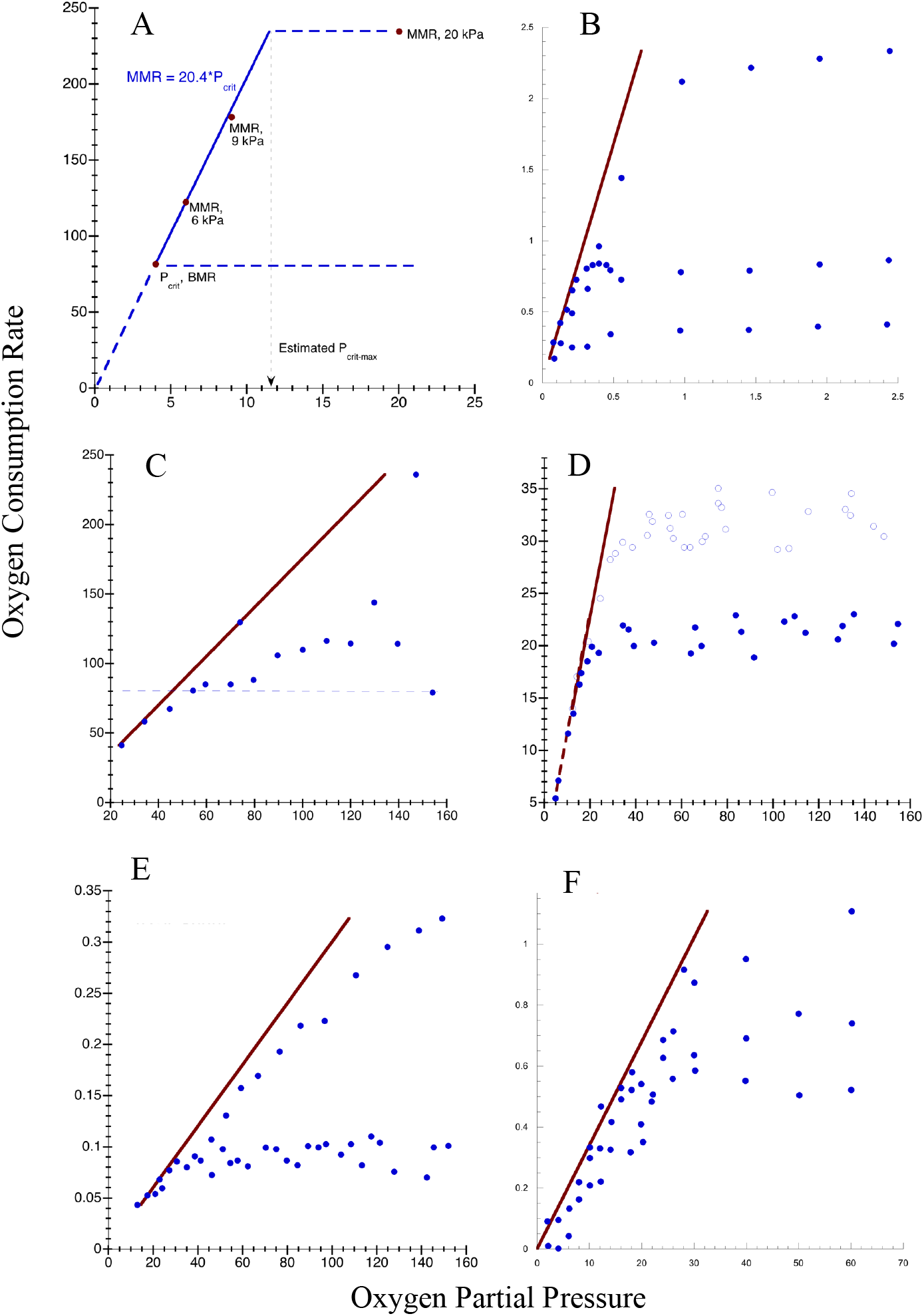
P_crit-*α*_ determined for the maximum achievable metabolic rate (MMR) at varying oxygen levels for seven aquatic species. A) The freshwater fish, *Lepomis gibbosus* (Crans et al., 2015): The data suggest that P_crit_ for MMR is less than air-saturation, meaning that the maximum rate recorded at 20 kPa could be maintained until ∼ 11.4 kPa PO_2_. B) *Gnathophausia ingens*, a deep-sea lophogastrid crustacean (Childress and Seibel, 1998). Three individuals measured at differing activity levels between maximum and rest. C) Lumpfish (*Cyclopterus lumpus*; Ern et al. 2016). D) Toadfish (*Opsanus tau*; Ultsch, 1989). Routine and standard metabolic rates. E) The prawn, *Penaeus esculentus* (Dall, 1986). F) *Octopus bimaculoides* (Seibel and Childress, 2000): Three individuals measured after varying acclimation periods resulting in variation in the metabolic rate. Units are as originally published.

Although rare, a continuous increase in *α*_*0*_ presents a challenge for P_crit-*α*_ determination because it is unclear whether oxygen supply had yet reached its capacity (i.e. whether *α*_*0*_ = *α*) or whether the final *α*_*0*_ represents measurement error. For most trials of this type, P_crit-*α*_ made visual sense and was not different from P_crit_ calculated using other methods (Fig. 4C). For a few others,*α* was unrealistically high, resulting in extremely low P_crit-*α*_. Some of these included positive MO_2_ data at zero PO_2_ or a positive y-intercept of the MO_2_ vs PO_2_ curve. In either case, this indicates continued oxygen consumption in the absence of oxygen. Because this is not physiologically possible, it must result from either PO_2_ error, perhaps introduced during sensor calibration, or inappropriate bin sizes. Both of these have drastic effects on the calculation of *α*, especially at low PO_2_ values where the ratio of error to true oxygen is magnified. If PO_2_ is underestimated (e.g. measurements include negative PO_2_ values due to a high zero calibration), *α*_*0*_ has the potential to diverge towards infinity as PO_2_ approaches zero. This occurs more often for species with very low P_crit_, such as those living in oxygen minimum zones and in stratified ponds. For example, using P_crit-*α*_, we could not support the finding of lower P_crit_ in goldfish following slow hypoxic induction as reported by Regand Richards (2017), but the majority of their trials achieved *α* at the lowest PO_2_ measured. Thus, we are less confident in our contrary finding.

We have developed a method to correct such PO_2_ error that will be described in full elsewhere (Timpe et al., in prep). It is included in the R package referenced above (Birk, 2020). Briefly, the correction involves identifying the magnitude of any PO_2_ error through incremental adjustments to PO_2_ values until two equivalent MO_2_ values are achieved for which *α*_*0*_ = *α*. This method effectively provides a way to diagnose and correct for calibration error in respirometry experiments. It is applicable only in specific circumstances and the only corrected data presented here are from the worm, *Sipunculus nudus* (Fig. 5). For this species, the correction resulted in *α* and P_crit-*α*_ values that more closely adhered to the MO_2_ vs PO_2_ curve.

## Discussion

We define Pcrit-_*α*_ as the PO_2_ at which physiological oxygen provision, *α*_*0*_, reaches its maximum capacity (*α*). Oxygen supply capacity, although species- and temperature-specific, is independent of PO_2_ or workload (i.e. metabolic rate). Thus, P_crit_ is a measure of *α* at any metabolic level from rest to maximum exertion and it is *α*, rather than P_crit_ *per se*, that contains relevant physiological information. Because *α* is not specific to a particular metabolic rate, P_crit-*α*_ need not be measured at SMR or any other specific metabolic rate. It is also not necessary for an individual animal to maintain a consistent metabolic rate throughout a trial or for an MO_2_ vs PO_2_ curve to have a clear break-point. P_crit-*α*_ can be calculated for any metabolic rate as long as oxygen supply reaches capacity at some point during the trial. P_crit-*α*_ is physiologically consistent and provides a simple, repeatable, and precise alternative to existing methods for P_crit_ determination. Moreover, this definition clarifies the physiological and evolutionary significance of P_crit_.

P_crit-*α*_ is a measure of the oxygen supply capacity. Selection for both elevated metabolic capacity (i.e. athleticism) and for metabolic performance in persistent hypoxia may elevate oxygen supply capacity (Fig. 8A). The P_crit-max_ (the PO_2_ above which no further increase in metabolic rate is possible) must be known to distinguish between these selective pressures (Fig. 8A). A P_crit-max_ less than air-saturation would suggest adaptation to persistent hypoxia. Most shallow-living aquatic or terrestrial species have a P_crit-max_ near atmospheric PO_2_ (21 kPa; Seibel and Deutsch, 2020). This means that additional environmental oxygen (hyperoxia) cannot elevate MMR. Among these “normoxic” species, most of the variation in P_crit_ reflects the differing temperature sensitivities of MMR and SMR and, thus, the factorial aerobic scope (FAS = MMR/SMR; Fig. 8B). Inserting MMR and SMR into *Eq. 1* and rearranging shows that FAS* P_crit_ = P_crit-max_. As such, the P_crit_ for normoxic species (P_crit-max_ ∼ 21 kPa) is inversely correlated with FAS (Seibel and Deutsch, 2020), explaining 95% of the variation (from 2-12 kPa) in P_crit_ among diverse species (Seibel and Deutsch, 2020). None of the variation could be attributed to measured differences in environmental PO_2_. Thus, P_crit_ alone, without also knowing MMR, is not a useful measure of hypoxia tolerance. Relative hypoxia tolerance in normoxic species must be conferred by adaptations for metabolic suppression and anaerobic metabolism that permit extended survival below P_crit_ (Mandic et al., 2009; Seibel, 2011).

**Figure 8.**
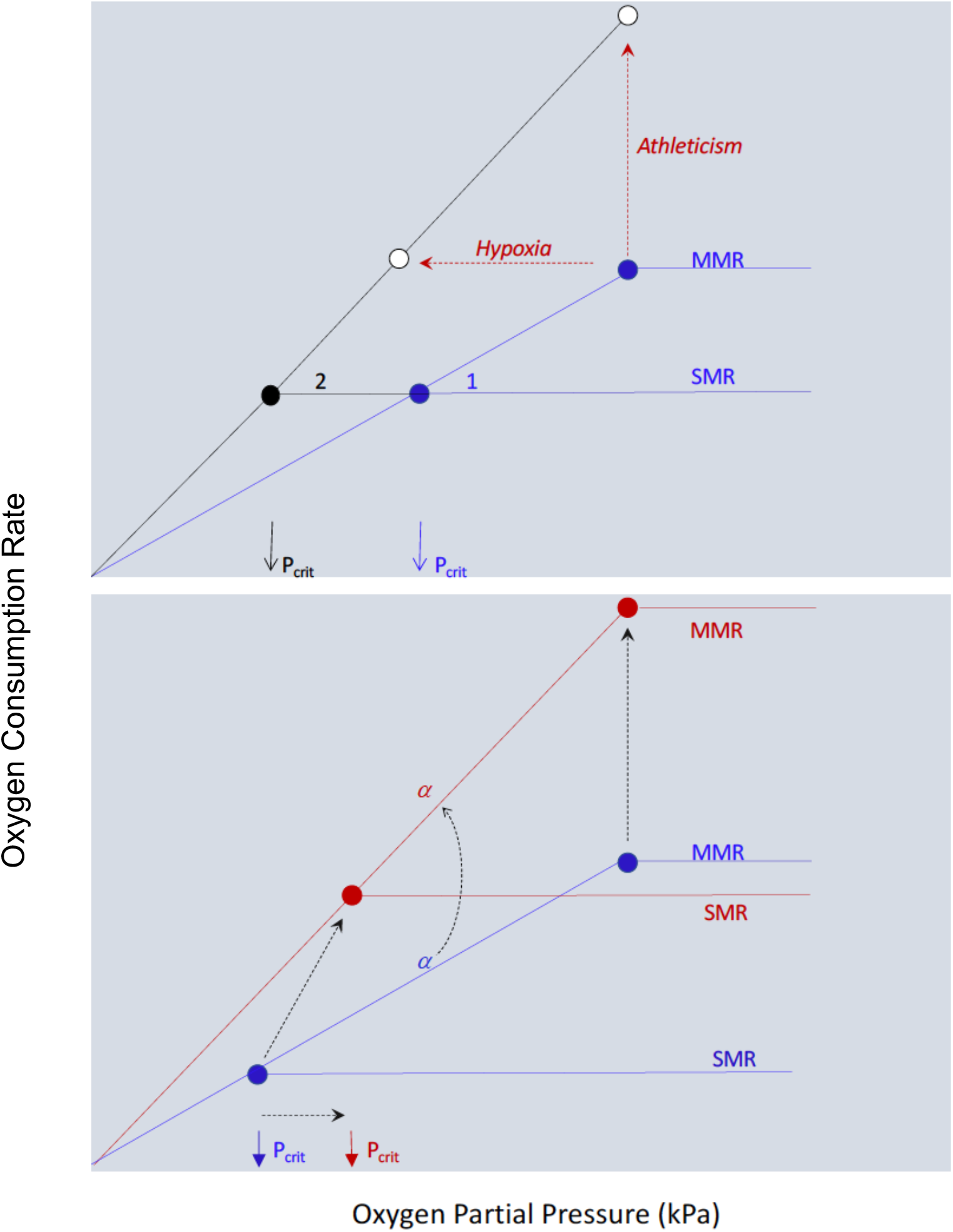
Schematics showing the relationships between SMR, MMR, P_crit_, oxygen supply capacity (*α*, slopes) and oxygen partial pressure. A) Two hypothetical species with identical SMR (1, blue; 2, black). Species 2 has a lower P_crit_ and, thus, a higher *α* (SMR/P_crit_; higher slope). Two possible MMR values (open circles) are shown for species 2. If both species have the same MMR, then P_crit-max_ (P_crit_ for MMR) must be lower for species 2, indicating hypoxia tolerance. Alternatively, the higher oxygen supply capacity may support a higher MMR and greater athleticism in normoxia for species 2. P_crit-max_ must be known to distinguish between these two alternatives. B) A single species depicted at two temperatures (warm, red; cold, blue). Black dashed arrows represent the increase in each metric (SMR, MMR and P_crit_) with temperature. The oxygen supply capacity increases with temperature to meet the higher MMR. SMR commonly increases with temperature faster than MMR (as shown here), causing P_crit_ to increase.

In contrast, adaptation to *persistent* hypoxia (that lasting longer than a diel or tidal cycle) results in a higher *α* for a given metabolic rate and P_crit_ is reduced at all metabolic levels, including MMR, relative to normoxic species. Thus, species that live in persistently hypoxic conditions have a lower P_crit_ *and* a lower P_crit-max_ relative to similar species in higher oxygen environments (Childress and Seibel, 1998; Seibel, 2011; Wishner et al., 2018; Mandic et al., 2009; Richards, 2011; Regan et al., 2019). Whether variation in P_crit_ and *α* reflects adaptation for increased metabolic capacity or hypoxia adaptation can only be known if MMR (and thus P_crit-max_) is also known (Fig. 8A). In the absence of such data, one can tentatively infer whether a high *α* is an adaptation for hypoxia or for metabolic capacity if it is assumed that FAS for the species of interest falls within the typical range (< ∼7; Seibel and Deutsch, 2020; Killen et al., 2016; Peterson et al., 1989). Thus *α* multiplied by 21 kPa (P_crit-max_ for normoxic species) should result in an estimated MMR less than ∼7X SMR. A greater value suggests that P_crit-max_ is less than 21 kPa by some degree and that oxygen supply mechanisms are adapted for hypoxia. Regardless, selection acts on the oxygen supply capacity in support of MMR and its P_crit-max_, not SMR and its P_crit_.

The maximum achievable metabolic rate (MMR) is linearly related to PO_2_ below P_crit-max_ and MMR reaches an oxygen limit at *α*, just as SMR does (Figs. 1, 7).For example, in the freshwater pumpkinseed fish, *Lepomis gibbosus* (Crans et al., 2015), measurements of P_crit_ at SMR and measurements of MMR at different oxygen levels show that the maximum achievable metabolic rate always becomes oxygen limited when PO_2_ = MR/*α*. The MMR measured in normoxia falls off the line, suggesting that *L. gibbosus* is hypoxia tolerant with a P_crit-max_ near 11 kPa (Fig. 7A). Similarly, *Gnathophausia ingens*, a deep-sea lophogastrid crustacean, maintained MMR down to ∼ 5 kPa while intermediate activity levels reached P_crit_ at proportionally lower PO_2_ equal to MR/*α* (Childress and Seibel, 1998). Distinct activity levels or metabolic states produce equivalent estimates of *α* for a variety of other animals including fishes (Ern et al., 2016; Ultsch et al., 1981), prawn (Dall, 1986) and octopus (Seibel and Childress, 2000; Fig. 7).

Both P_crit_ and SMR typically increase with temperature (Rogers et al., 2016 for review), although the temperature sensitivities (*E*, eV) for each metric are quite different. The typically lower temperature sensitivity of Pcrit, relative to SMR, has led some to suggest that oxygen supply does not keep pace with oxygen demand (Pörtner et al., 2017; Keilland et al., 2019; Deutsch et al., 2020). However, this makes little sense given that oxygen supply is sufficient to meet MMR, which is typically many times higher. In contrast, oxygen supply matches MMR with increasing temperature. If MMR increased with temperature as fast as SMR, P_crit_ would not change (Fig. 8B). Mathematically, the *E* value (exponent) for *α* must equal the difference between the *E* values for SMR and P_crit_. If SMR increases faster with temperature than P_crit_, *E*_*α*_ must be less than *E*_SMR_, leading to the assumption that oxygen supply does not match demand at high temperatures. However, since P_crit_ for MMR (P_crit-max_) is effectively the maximum environmental PO_2_, it does not change with temperature for most species (*E*_Pcrit-max_ = 0). Thus *E*_*α*_ exactly matches *E*_MMR_ and oxygen supply increases with temperature to match maximum demand (Fig. 6A). For example, in the lumpfish, the measured *E*_*α*_ (0.29 eV), derived from SMR and P_crit_, is very similar to the measured *E*_MMR_ (0.27 eV). The fact that P_crit_ and *α* do not increase with temperature in proportion to SMR *in no way* indicates oxygen limitation at high temperature. To the contrary, oxygen supply evolved to meet maximum demand regardless of temperature.

The definition of P_crit-*α*_, provided here, requires reassessment of the concept of oxygen consumption regulation (oxyregulation). Regulation identifies the ability of a species to increase oxygen provision as oxygen availability declines. However, a variable or declining metabolic rate during a respirometry experiment may merely reflect behavioral changes or changes in metabolic state unrelated to oxygen and is not necessarily diagnostic of a physiological inability to regulate. Oxyconformation (a continuous decline in MO_2_ throughout a respirometry trial), in contrast, implies that an individual is operating at its maximum oxygen supply capacity throughout a trial regardless of PO_2_. Accordingly, it suggests that either the individual is operating at the maximum achievable metabolic rate for a given PO_2_ or that it completely lacks aerobic scope (i.e. MMR equals SMR regardless of oxygen availability). For conformation to occur, physiological oxygen supply mechanisms must be either absent, always operating at their maximum capacity, or incapable of regulating their output. Thus, true conformation can be diagnosed by a constant oxygen supply (MR/PO_2_) as PO_2_ declines (Gnaiger, 1993). For example, a reanalysis of a representative respirometry trial for *Sipunculus nudus*, a worm that is widely regarded as an oxyconformer (Pörtner and Grieshaber, 1993), shows that *α* increases as PO_2_ declines (Fig. 5), suggesting some level of active regulation, or what Gnaiger (1993) referred to as microoxic regulation. Oxyconformation is seemingly rare (Ultsch and Regan, 2019), even among animals that lack complex circulatory systems (Rutherford and Thuesen, 2005).

The determination of *α*, rather than P_crit_ alone, not only alters our perception of the ecological and evolutionary significance of P_crit_, it is also a refinement of the respirometry methodology required to determine P_crit_. Typically, P_crit_ and SMR are determined during the same trials, requiring a lengthy acclimation period to ensure a resting, fasted state. Not only are such experiments extremely time-consuming, but an acclimatory response in the oxygen supply system may influence P_crit_, especially in fishes with highly elastic physiologies. For example, goldfish are known to remodel gills during hypoxic exposure, which reduces P_crit_ (Sollid et al., 2005; Regan and Richards, 2017). The present method eliminates the requirements for long experiments and for slow hypoxic induction to determine P_crit_. For example, trials using a conventional broken stick method must extend to oxygen levels well below P_crit_ solely to produce sufficient data points for a reliable regression of the oxygen dependent portion of the curve. In contrast, P_crit-*α*_ trials can be truncated by starting at relatively low PO_2_ and/or by measuring relatively high (active) metabolic rates where oxygen supply is maximized above potentially lethal oxygen levels. P_crit-*α*_ trials can be terminated as soon as *α* has plateaued or reached a summit, thus avoiding potential distress in experimental animals in hypoxic conditions just to satisfy statistical requirements for data below P_crit_. An additional benefit of this method is that once *α* has been determined for a given species and temperature, P_crit-*α*_ can be calculated for any physiological state or metabolic rate of interest (e.g. following feeding or exercise) simply by measuring their respective metabolic rates in normoxia. This facilitates modeling of habitable environments based on the metabolic responses to a variety of environmental and physiological conditions (Deutsch et al., 2020; 2015).

## Conclusions

The critical oxygen pressure is redefined here (P_crit-*α*_) as the PO_2_ at which the oxygen transport machinery reaches maximum capacity. The advantages of this definition include 1) P_crit-*α*_ is directly measured without the need for complex statistics that hinder measurement and interpretation; 2) it makes clear that P_crit_ is a measure of oxygen supply, which does not necessarily reflect hypoxia tolerance; 3) it alleviates many of the methodological constraints inherent in existing methods; 4) it provides a means of predicting the maximum metabolic rate achievable at any PO_2_, 5) P_crit-*α*_ sheds light on the temperature- and size-dependence of oxygen supply and metabolic rate and 6) P_crit-*α*_ can be determined with greater precision than traditional P_crit_. We argue that P_crit-*α*_ provides important physiological information that is lost in non-linear analyses such as the *Regulation Index*. Moreover, we argue that the concept of oxyregulation should be redefined with reference to the physiological adjustments that increase oxygen delivery as environmental PO_2_ declines, rather than the maintenance of an arbitrary metabolic rate. P_crit-*α*_ provides greater predictive power and enables testing of previously obscured hypotheses regarding metabolic scope.

## Acknowledgements

This project was supported by NOAA grants NA18NOS4780167 and NA17OAR4310081 and NSF grant OCE-1459243 to BAS., the Jack and Katharine Ann Lake Fellowship to AA, the Anne and Werner Von Rosenstiel and Garrels Memorial Endowed Fellowships to AT, The Hogarth Fellowship to CW, The Southern Kingfish Association Fellowship to AB, and a National Science Foundation postdoctoral fellowship (DBI-1907197) to MAB.

## Data Availability

Citations are provided for all published data used in this paper. Unpublished datasets are summarized in Table 2 and full datasets will be made available upon publication of their primary sources.

